# Protein restriction associated with high fat induces metabolic dysregulation without obesity in juvenile mice

**DOI:** 10.1101/2024.07.22.604540

**Authors:** Amélie Joly, Jean-Louis Thoumas, Anne Lambert, Estelle Caillon, François Leulier, Filipe De Vadder

## Abstract

Dysregulation of energy metabolism, including hyperglycemia, insulin resistance and fatty liver have been reported in a substantial proportion of lean children. However, non-obese murine models recapitulating these features are lacking to study the mechanisms underlying the development of metabolic dysregulations in lean children. Here, we develop a model of diet-induced metabolic dysfunction without obesity in juvenile mice by feeding male and female mice a diet reflecting Western nutritional intake combined with protein restriction (mWD) during 5 weeks after weaning. mWD-fed mice (33% fat, 8% protein) do not exhibit significant weight gain and have moderate increase in adiposity compared to control mice (16% fat, 20% protein). After 3 weeks of mWD, juvenile mice have impaired glucose metabolism including hyperglycemia, insulin resistance and glucose intolerance. mWD also triggers hepatic metabolism alterations, as shown by the development of simple liver steatosis. Both male and female mice fed with mWD displayed metabolic dysregulation, which a probiotic treatment with *Lactiplantibacillus plantarum* WJL failed to improve. Overall, mWD-fed mice appear to be a good preclinical model to study the development of diet-induced metabolic dysfunction without obesity in juveniles.

## Introduction

Childhood represents a critical developmental window during which environmental cues, notably nutrition, significantly influence body mass index and energy metabolism [1]. Notably, pediatric overweight and obesity result from an energy intake exceeding a child’s requirement, and affects almost 40 million children under 5 worldwide [2]. It is estimated that 6-39% of obese children will develop metabolic dysregulation, including elevated blood glucose, insulin resistance and metabolic dysfunction-associated steatotic liver disease (MASLD), which elevate the risk of metabolic disorders such as type 2 diabetes and cardiovascular diseases [3,4]. Even though its prevalence is much higher in obese and overweight children, metabolic dysregulation can also occur independently of BMI increase in children [5–9]. Indeed, a recent retrospective study has shown that MASLD may affect 8% of lean American adolescents [10]. Likewise, the prevalence of insulin resistance, low HDL cholesterol and elevated plasma glucose among non-overweight and non-obese children can reach 2.2%, 8.1% and 6.6% respectively [11]. However, the mechanisms underlying the development of these metabolic alterations in lean children are poorly understood.

Rodent models recapitulating the features of metabolic dysfunction without obesity are key for comprehending these mechanisms and their short and long-term effects on organismal physiology. However, current rodent models do not capture the diverse spectrum of metabolic dysfunctions observed in humans, nor accurately reflect the nutritional context under which these metabolic alterations develop in lean children. Specifically, diets used to model pediatric metabolic dysfunction without obesity are often extremely deficient in specific amino acids [12–14] or contain levels of fats or sugars that are too high to reflect the average intake in a Western diet [15,16]. Moreover, these murine models have been extensively described in adults, but to date very few rodent studies have investigated their effects in juvenile animals [17].

In this study, we aimed to bridge this gap by developing a model of metabolic dysfunction without obesity in juvenile mice, with a nutritional challenge consistent with the dietary intakes of children. To this end, we adapted the new total western diet (ntWD), which is a rodent diet designed to reflect dietary intake of overweight Americans and contains physiological amounts of fat and sugars from different sources [18]. Previous work has shown that the ntWD induces substantial weight gain, but not obesity in adult mice, without clear modification of glucose metabolism [18,19]. Building on the fact that dietary amino acid restriction in juvenile mice is associated with glucose metabolism alterations [20–22], as well as hepatic steatosis onset [23,24], we designed a modified version of the ntWD with a reduced amount of protein that we called the modified western diet (mWD). In juvenile mice, this diet induced features of a metabolic dysregulation including hyperglycemia, insulin resistance and hepatic steatosis, without substantial body mass index increase. The mWD therefore provides a model of diet-induced metabolic dysfunction without obesity in juvenile mice.

## Material and methods

### Animals and diet

Animal experiments have been carried out in accordance with the European Community Council Directive of September 22^nd^, 2010 (2010/63/EU) regarding the protection of animals used for scientific purposes. All experiments have been approved by a local ethic committee (CECCAPP) and by the French Ministry of Research (APAFIS #40481-2022103017295302 v3).

Adult conventional C57Bl/6N male and female mice were purchased from Charles River. After one week of acclimatation, animals were mated and their offspring was used for experiments. The litters were reduced to 6 pups per dam at post-natal day 2. At post-natal day 21, male or female offspring were randomly weaned on control AIN93G diet containing 63.6% of carbohydrates, 15.9% of lipids and 20.5% of proteins (CD) or on the experimental diet corresponding to a modified version of the new total Western Diet [18] with moderate protein restriction and containing 58.3% of carbohydrates, 33.4% of lipids and 8.3% of proteins (mWD). Details of diet composition are provided in Table S1. Animals were weighed and measured once a week after flash anesthesia with 3% isoflurane, from weaning until post-natal day 56. Food intake was calculated on a weekly basis by subtracting the amount of remaining food from the amount provided the week before. At post-natal day 56, animals were fasted during 6h starting at 8 AM, flash anesthetized with 3% isoflurane and killed by cervical dislocation. Liver, perigonadal adipose tissue and subcutaneous adipose tissue were weighed and liver was kept in PBS-30% sucrose to be processed for histology or snap-frozen in liquid nitrogen and kept at −80°C until further analysis.

#### Probiotic treatment

Mice were treated 5 times a week with 10^9^ CFU/100 µL of *Lactiplantibacillus plantarum* WJL, or a placebo solution containing maltodextrin, administered using a pipette in the mouth of the mouse. Animals were held until they swallowed the solution.

### Glucose metabolism assessment

An oral glucose tolerance test was performed at post-natal day 42, after 3 weeks of feeding with experimental diets. Animals were fasted during 6h starting at 8 AM and then gavaged with 2 mg/g of D-(+)-glucose (Sigma Aldrich, G70021). Glycemia was measured from the tail vein using a Verio Reflect glucometer (OneTouch) before (0 min) and 15, 30, 45, 60, 90 and 120 min after glucose gavage. In addition, blood samples were collected from the tip of the tail vein at 0, 15 and 30 min using Microvette© CB 300 Z (Starstedt) and centrifuged 5 min at 10,000 g. Serum was stored at −20°C until insulin measurement using Ultra Sensitive Mouse Insulin ELISA kit (Crystal Chem, 90080), following the manufacturer’s instructions.

### Lipid metabolism assessment

Liver triglycerides were measured using the GPO method (Biolabo, 87319) according to the manufacturer’s instructions.

Liver sections were stained with Oil Red O as previously described [24]. Briefly, liver samples were dehydrated overnight in 30% sucrose (Roth, 4621.1), embedded in OCT (Leica, 14020108926) and kept at −80°C until further processing. 10 µm-thick sections were cut using cryostat (Leica, CM 3050S), fixed in 4% paraformaldehyde (Sigma, P6148) and stained with Oil Red O (Sigma, O-0625). Sections were imaged using a DM6000 microscope (Leica) at 20x magnification. 10 images were taken per mouse, on 4 different slides, choosing random fields on each slide. Images were analyzed with Fiji software [25], using a modified version of the macro developed by Nishad et al. [26], with the following parameters: threshold (0 – 95) and particle size (10 – 250 µm^2^).

### RNA extraction and RT-qPCR

Total RNA from liver samples was extracted and purified using the Nucleospin RNA kit (Macherey-Nagel, 740955.250). RNA quality and quantity was assessed using Nano Drop 2000 (Thermo Fisher Scientific) and 1 µg of RNA was retrotranscribed with the Sensifast cDNA synthesis kit (Meridian Biosciences, BIO-65053). Quantitative PCR was performed on a Bio-rad CFX96 apparatus using Takyon^TM^ No ROX SYBR Mastermix blue dTTP (Eurogentec, UF-NSMT-B0701), 1/5 cDNA dilutions of the reverse transcription product and specific primer sets (Table S2). Analysis was performed according the MIQE guidelines [27]. Amplification efficiency was calculated for each primer set using cDNA serial dilutions and melting curves for amplicons were analyzed to ensure unique and specific amplification. *Tbp*, *Rpl32* and *Actb* were used as control genes after confirming their stability using the geNorm tool available in the CFX Manager^TM^ software (Biorad). Data are expressed as geometric mean relative to the expression of control genes.

### Statistical analysis

Statistical analysis was performed with GraphPad Prism software version 10.2.1 (GraphPad software, San Diego, California, USA, www.graphpad.com) and using parametric Student’s t-test or the non-parametric Mann-Whitney test. Male and Female datasets were analyzed separately to assess specifically the effect of the diet in each sex without the interaction between the two parameters. Data were considered to show evidence for an effect when the p-value was inferior to 0.05.

## Results

### 1. mWD does not induce obesity in juvenile mice

To explore the effects of modified western diet (mWD) feeding on physiology of juvenile mice, we fed mice with mWD and as control CD diet for 5 weeks. Both male and female mice fed with mWD during 5 weeks after weaning did not develop obesity or overweight (Fig 1 A-B), contrary to what has been observed in response to classic high-fat feeding [28–30]. Indeed, males displayed similar weight-for-length ratio compared to control diet-fed mice (Fig 1C-D). mWD-fed females gained slightly more weight than CD animals, since they weighed on average 1.2 g more than controls at post-natal day 56 (Fig 1B). In line with that, analysis of covariance (ANCOVA) revealed that mWD females had slightly higher weight-for-length ratio than CD females (Fig 1D). Despite a modest weight increase, we can consider that mWD fed females did not develop overweight or obesity, similar to males. Consistently, mWD did not induce changes in subcutaneous fat ratio (Fig 1E, H). However, juvenile male but not female mice fed a mWD seemed to accumulate more visceral adipose tissue, evaluated by the mass of perigonadal fat (Fig 1F, I), a type of visceral fat, which is a typical feature of metabolic dysfunction [31]. Interestingly, mWD-fed males displayed decreased muscle mass, evaluated by the weight of the gastrocnemius muscle, compared to CD animals (Fig 1G). This decrease in muscle mass might explain why they have no change in total body weight despite increased VAT accumulation. On the contrary, mWD-fed females exhibited no change in muscle mass (Fig 1J).

**Figure 1:**
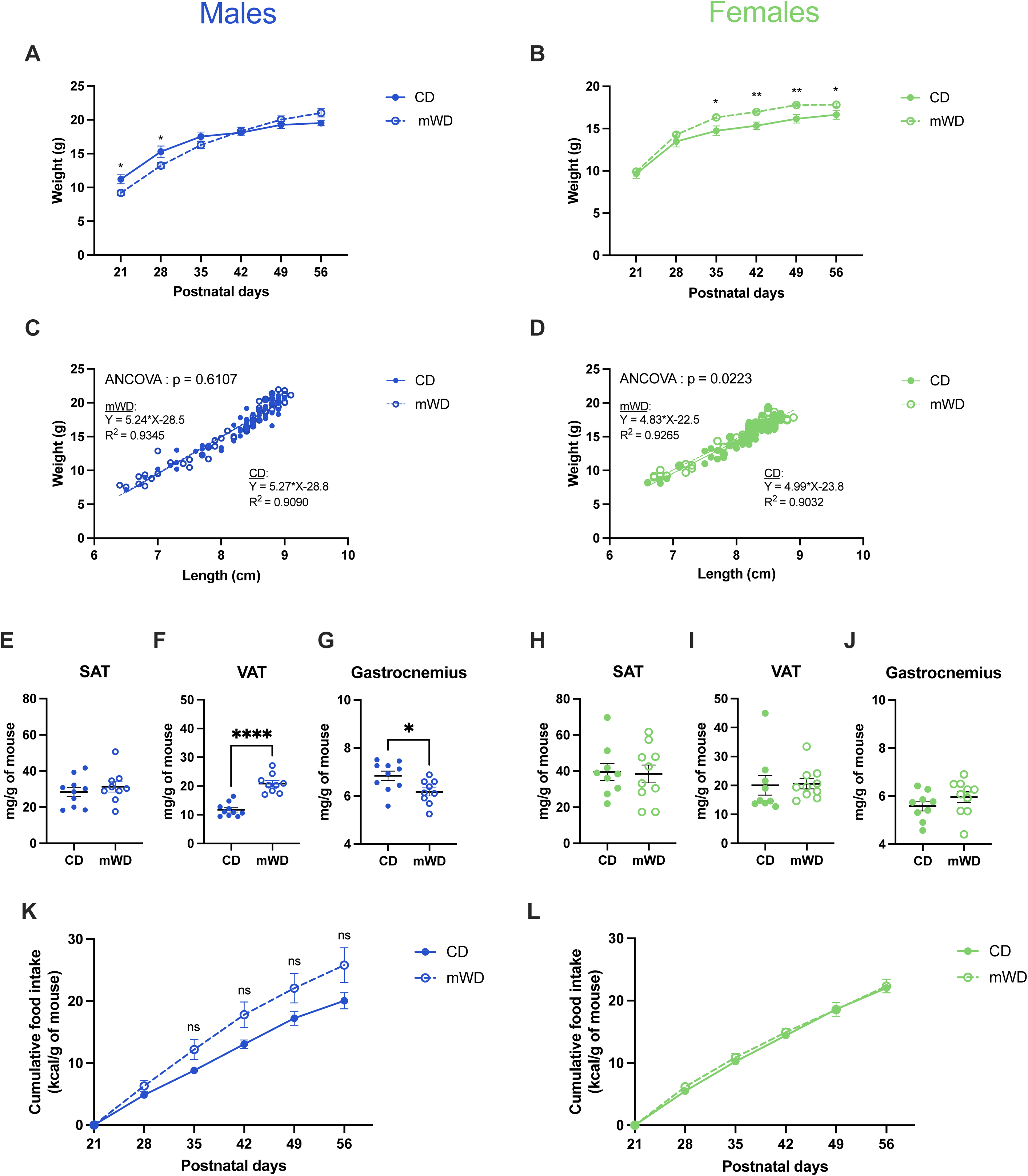
Modified western diet feeding to juvenile male and female mice does not induce obesity. (A-B) Body weight of male and female mice fed control diet (CD) or modified western diet (mWD) between weaning (postnatal day 21, P21) and P56. (C-D) Length-versus-weight of CD- and mWD-fed males and females. (E-J) Weight of subcutaneous adipose tissue (SAT), visceral adipose tissue (VAT) and gastrocnemius muscle measured at P56 and expressed per gram of body weight. (K-L) Cumulative food intake. Data are expressed as mean ± SEM. * p<0.05, ** p<0.01, **** p<0.0001. (A-J): unpaired t-test. (K-L): Mann-Whitney test.

The lack of weight gain in mWD fed animals was not due to a modification of caloric intake, as cumulative food intake was similar between mWD and CD animals (Fig 1K-L). However, this seemingly similar caloric intake hid discrepancies in major nutrient intake, since mWD-fed mice consumed twice as much fat and half as much protein as control mice (Figure S1). Therefore, mWD during the juvenile period is not obesogenic despite increased accumulation of visceral fat in a sex-dependent manner.

### 2. mWD triggers glucose metabolism alterations that are independent of sex

We then characterized glucose metabolism in juvenile male and female mice after 3 weeks of mWD or CD feeding. Both males and females developed hyperglycemia with respectively 38% and 47% increase in fasting glycemia compared to controls (Figure 2 A, F). However, mWD feeding did not trigger changes in fasting insulinemia (Figure 2B, G). In accordance, homeostatic model assessment of insulin resistance (HOMA-IR) calculation showed evidence for insulin resistance in these animals (Figure 2C, H). Thus, mWD feeding induces alterations of glucose metabolism during fasting. An oral glucose tolerance test demonstrated that both mWD-fed males and females had impaired glucose tolerance (Figure 2D, E, I, J). Altogether, these data indicate that mWD feeding during the juvenile period triggers a complete dysregulation of glucose homeostasis, independently of any weight gain and obesity.

**Figure 2:**
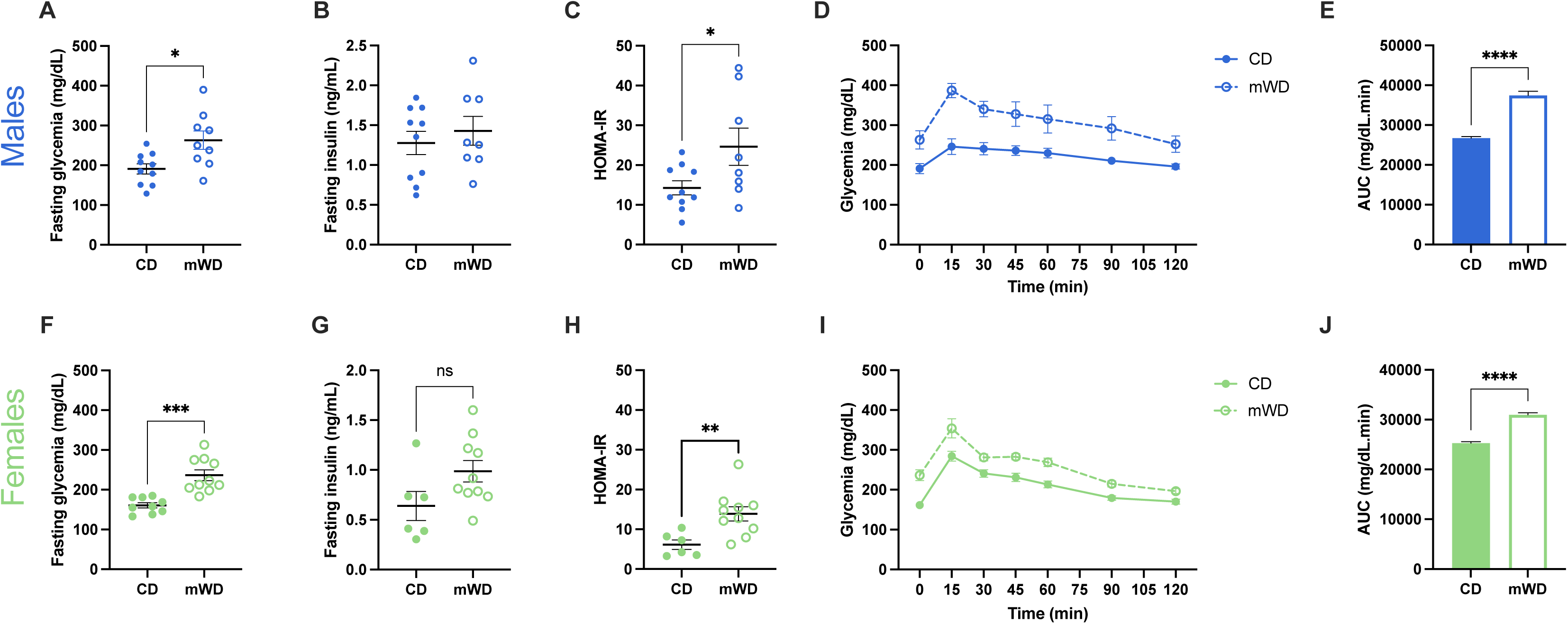
Male and female mice fed mWD after weaning have impaired glucose metabolism. Fasting glycemia (A, F) and insulinemia (B, G) of control (CD) or mWD-fed males and females at post-natal day 42 (P42). (C, H) Homeostatic model assessment of insulin resistance (HOMA-IR) calculated at P42. (D, I) Oral glucose tolerance test performed on CD and mWD males and females at P42 and area under the curve for this test (E, J). Data are expressed as mean ± SEM. * p<0.05, ** p<0.01, *** p<0.001, **** p<0.0001, unpaired t-test.

### 3. mWD feeding induces simple hepatic steatosis development

In addition to glucose metabolism dysfunction, lean children and adolescent can develop MASLD [10]. We thus investigated the extent to which mWD altered liver metabolism. mWD-fed animals did not develop hepatomegaly (Figure 3A, C). However, significant liver triglyceride accumulation was detected in response to juvenile mWD feeding (Figure 3B, D), indicating that these animals suffer from liver steatosis. Consistently, Oil Red O staining revealed increased lipid droplet size upon mWD (Figure 3E-F).

**Figure 3:**
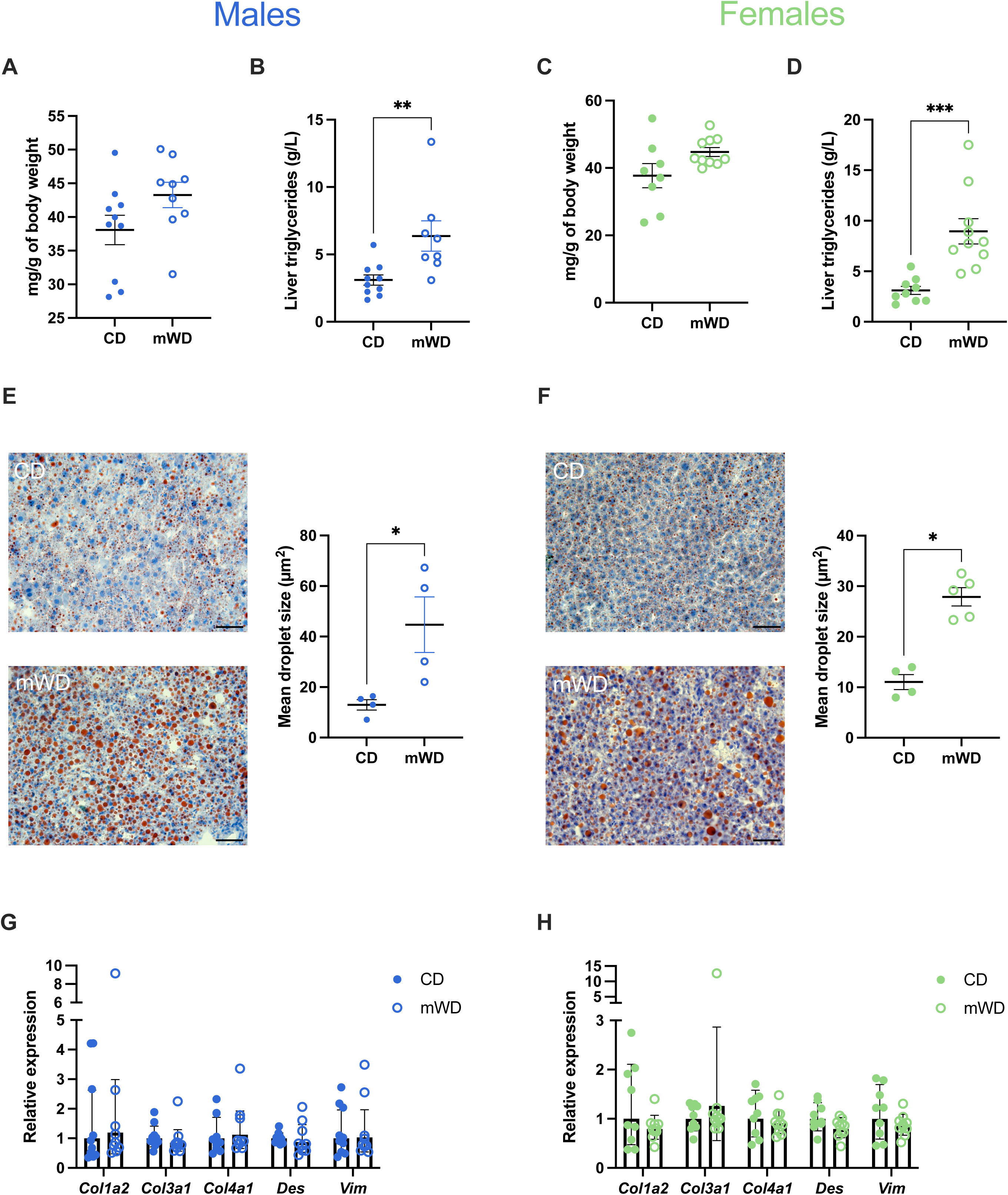
mWD feeding during the juvenile period triggers simple hepatic steatosis in both males and females. Weight (A, C) and triglyceride concentration (B,D) in the liver of CD- and mWD-fed males (blue) and females (green) at post-natal day 56 (P56). (E, F) Representative images of Oil Red O stained liver sections and quantification of mean lipid droplet sizes. Scale bars: 50 µm. (G, H) Liver *Col1a2*, *Col3a1*, *Col4a1*, *Des* and *Vim* expression in CD- and mWD-fed males and females. (A-F): Data are expressed as mean ± SEM. (G, H): Data are expressed as geometric mean ± geometric SD. * p<0.05, ** p<0.01, *** p<0.001. (B, D): unpaired t-test. (E, F): Mann-Whitney test.

Dyslipidemia, defined as elevated circulating levels of triglycerides or decreased levels of high-density lipoprotein [32] is included in the scope of symptoms associated to metabolic dysfunction. Circulating triglyceride levels were similar in the portal vein and cava vein of mWD and CD fed animals (Figure S2A, B, D, E), providing no evidence for dyslipidemia development in our model. These data suggest no increased triglyceride absorption from the gut nor modified liver triglyceride export. This is consistent with previous findings supporting the idea that liver steatosis is associated with *de novo* lipid synthesis in the liver [33]. However, we did not detect modified expression of *Fasn* and *Acc2*, two enzymes involved in triglyceride synthesis (Figure S2C, F). mWD-fed females had decreased hepatic expression of *Cpt1*, a rate-limiting enzyme involved in the beta oxidation of long-chain fatty acids, suggesting that lipid catabolism could be altered upon mWD in females. However, no change in the expression of another enzyme involved in lipid oxidation, *Acaa2*, was detected (Figure S2F). In males, mWD feeding was associated with a two-fold reduction in hepatic *Hmgcr* expression, suggestive of decreased cholesterol synthesis in the liver of these animals. Surprisingly, the expression of *Acc2,* which is a rate-limiting enzyme for fatty acid synthesis, was also decreased in the liver of mWD-fed males compared to CD males. This decrease in *Acc2* expression contrasts with the expected increase in fatty acid synthesis in the steatotic liver. These findings suggest that mWD feeding induces complex alterations in hepatic lipid metabolism, potentially involving compensatory mechanisms. In addition, no change in the expression of genes encoding proteins involved in lipid transport (*Fabp1*) or in very-low density lipoprotein synthesis (*Mttp*) or export (*Surf4*) was detected.

MASLD is a disease that evolves throughout time from simple steatosis to liver fibrosis, cirrhosis and hepatocellular carcinoma [34]. Hepatic fibrosis is associated with the appearance of several histological features including extracellular matrix development [34]. To test whether mWD feeding induced liver fibrosis in addition to steatosis, we looked for markers of extracellular matrix accumulation. We detected no increased expression of several genes encoding collagen subunits (*Col1a2*, *Col3a1*, *Col4a1*) nor of genes coding for the extracellular matrix components desmotubulin (*Des*) and vimentin (*Vim*,) in the liver of male and female mice fed mWD (Figures 3G, H). Therefore, 5 weeks of mWD feeding during the juvenile period appears to lead to the development of simple liver steatosis, but not to fibrosis.

### 4. A chronic treatment with the probiotic bacterium LpWJL fails to rescue metabolic dysfunction induced by mWD

Modulating the gut microbiome through probiotic supplementation represents a promising therapeutic approach that has been shown to favorably impact metabolic parameters [35]. In particular, we previously reported the beneficial impact of the the probiotic bacterium *Lactiplantibacillus plantarum^WJL^* (LpWJL) on glucose homeostasis of juvenile mice fed a low-protein low-fat diet [20,36]. In particular, chronic LpWJL treatment partially rescued insulin production and glucose tolerance in malnourished animals [20]. We thus hypothesized that LpWJL could ameliorate the features of metabolic dysfunction triggered by another nutritional challenge during the juvenile period: mWD feeding. We chronically treated juvenile mWD-fed mice with 10^8^ CFU of LpWJL or with a placebo during 5 weeks after weaning. LpWJL did not induce modification of body weight and subcutaneous adipose tissue proportion in males and females (Fig 4A-B, G-H). Interestingly, male, but not female, mice treated with LpWJL displayed decreased visceral adipose tissue ratio (Fig 4C, I), a feature that was specifically altered by mWD in males (Fig 1F). However, LpWJL treatment failed to rescue glucose metabolism and hepatic alterations induced by mWD (Figure 4D-F, J-L). These data demonstrate that the beneficial effect of LpWJL treatment on the juvenile host is highly dependent on the nutritional context, with a positive impact upon chronic protein and lipid malnutrition but not upon mWD [20]. Future investigations will have to evaluate the ability of other specific probiotic strains or consortia to ameliorate metabolic abnormalities in non-obese juvenile mice.

**Figure 4:**
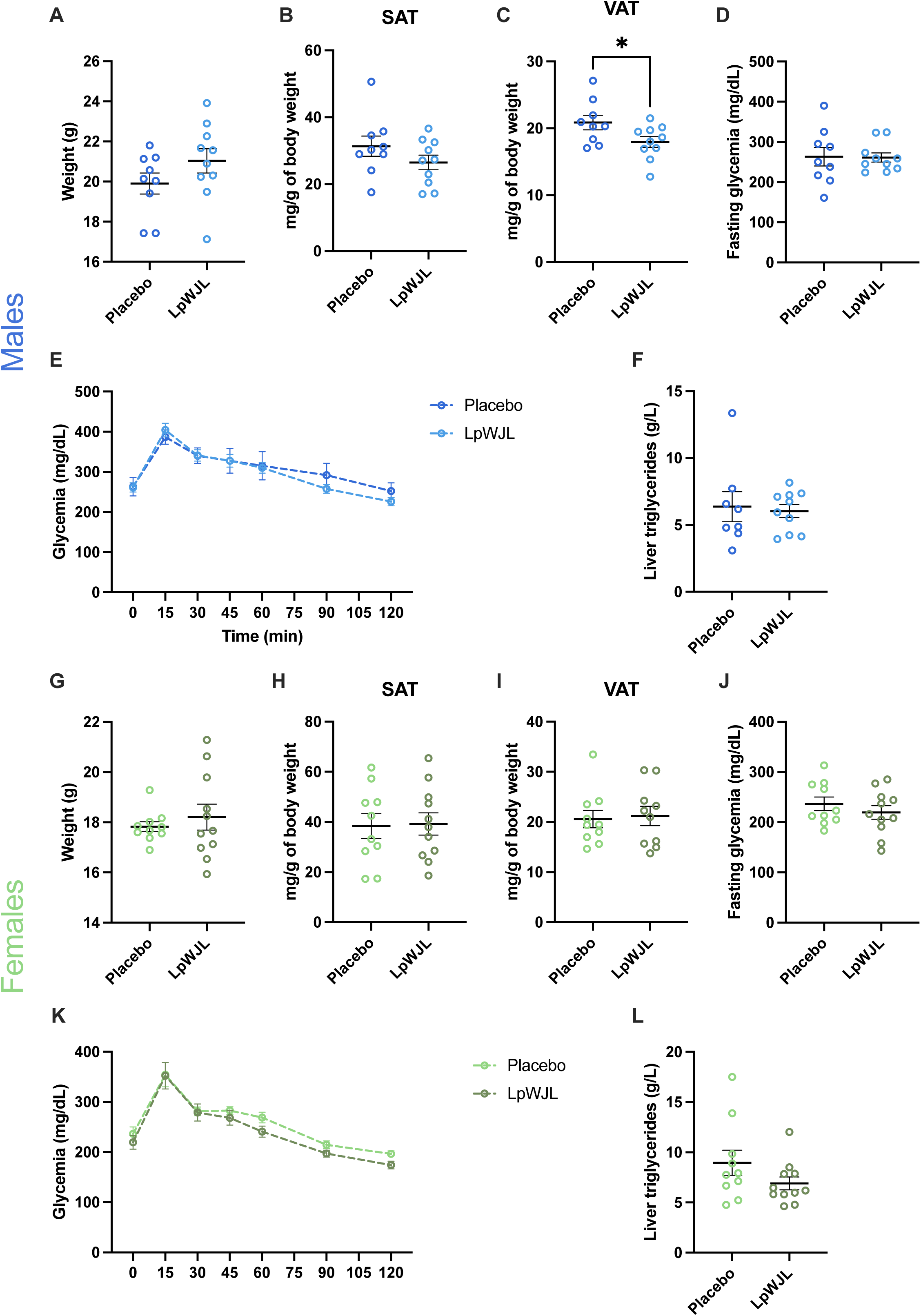
LpWJL treatment fails to improve metabolic dysfunction induced by mWD feeding during the juvenile period. Weight of body (A, G), subcutaneous adipose tissue (B, H) and visceral adipose tissue (C, I) of male (blue) and female (green) mice fed mWD and treated with placebo or with LpWJL at post-natal day 56. (D, J) Plasma glucose after 6 hours of food deprivation and oral glucose tolerance test (E, K) performed at post-natal day 42. (F, L) Liver triglyceride content. Data are expressed as mean ± SEM. * p<0.05, unpaired t-test.

## Discussion

The prevalence of children suffering from metabolic dysfunction without obesity is non-negligible and these children have an increased risk of mortality and development of metabolic diseases [5,9]. Thus, the establishment of rodent models recapitulating the features of this disorder is crucial to study the mechanistic basis of its onset and evolution. In this study, we describe a modified version of the new total western diet [18] with moderate protein restriction, which triggers the development of metabolic dysfunction in juvenile mice without inducing obesity.

Juvenile male and female mice fed a mWD during 5 weeks did not develop obesity, contrary to what has been described in this strain of mouse in response to feeding with a high fat diet during the juvenile period [28–30]. However, mWD-fed animals had metabolic dysfunction, including hyperglycemia, impaired glucose tolerance, signs of insulin resistance and simple liver steatosis, but did not present evidence of liver fibrosis. In murine models, fibrosis usually takes several months to develop. For example, it was observed after 8 weeks of high-fat high-sucrose diet in a mouse model of juvenile MASLD [17]. In this study, we wanted to address metabolic defects induced by mWD during the juvenile and peri-weaning period. We could hypothesize that hepatic fibrosis would develop upon longer feeding with mWD. Of note, it is evaluated that approximately 25% of fatty livers will progress toward steatosis and cirrhosis, the other 75% remaining simple steatosis [37]. Therefore, the simple steatosis that we capture in our model is representative of the majority of cases. The molecular mechanisms underpinning the development of hepatic steatosis upon mWD feeding remain to be clarified. Our analysis did not reveal notable alterations in specific hepatic lipid metabolism pathways based on the expression of selected genes. However, we identified signatures suggesting that mWD-fed males exhibit decreased hepatic cholesterol synthesis, while lipid oxidation might be altered in the liver of mWD-fed females. In line with these hypotheses, alterations in these processes have been observed in the liver of juvenile rodents subjected to a low-protein diet [23,24,38].

Metabolic syndrome (MetS) constitutes a cluster of symptoms, including elevated blood glucose, insulin resistance, MASLD, hypertension and obesity, which elevate the risk of metabolic disorders such as type 2 diabetes and cardiovascular diseases [4]. However, the criteria for MetS proposed by different organizations have not been fully harmonized [39]. The type of metabolic dysfunction that we describe in this study, occurring independently of some typical MetS symptoms, highlights that metabolic disturbances can manifest in different ways, and that the traditional criteria for MetS may not fully capture the heterogeneity of metabolic dysfunction, especially in lean populations. Overall, mWD induces metabolic defects that recapitulate many features of MetS in juveniles, but further work will be required to fully characterize this model, and in particular its relevance for the cardiovascular disorders associated with MetS.

Consumption of a diet presenting a western-type nutrient profile by adult male mice does not induce metabolic disorder nor weight gain [18]. Here, we show that modifying this nutrient profile by decreasing the amount of protein is sufficient to induce metabolic dysfunction in male and female juvenile mice. Low-protein feeding has been associated with an increased energy expenditure in several rodent studies [24,40], even though other studies evidenced no effect of protein malnutrition on energy expenditure [21,41]. Thus, we cannot exclude that a reduction in dietary protein intake could contribute to an increased energy expenditure preventing obesity in our model. Interestingly, we recently showed that male mice fed mWD between 35 and 119 post-natal days did not develop obesity, hyperglycemia nor glucose intolerance [19], highlighting that the post-weaning period is key for the development of metabolic alterations.

Several other rodent models of diet-induced metabolic dysfunction without obesity have been previously described. However, they are associated with nutrient composition that are not physiologically relevant to human health and usually induce weight loss. Mice fed methionine- and choline-deficient diets have impaired phosphatidylcholine synthesis and VLDL assembly, which causes hepatic triglyceride accumulation and MASLD [13,14]. However, this diet induces significant body weight loss [12,13] and does not trigger insulin resistance [14]. Besides, choline-deficient amino acid defined diets mimic histological features of MASLD and avoid extreme weight loss but do not induce insulin resistance nor impaired glucose tolerance [13]. To our knowledge, our model is the first to induce metabolic dysfunction without obesity in juvenile mice, using a diet that does not involve extreme deficiencies or excesses in essential nutrients.

Metabolic dysfunctions often are sexually dimorphic. For example, men are at higher risk of developing more severe liver fibrosis compared to pre-menopausal but not post-menopausal women [42]. In addition, females are less susceptible to develop metabolic dysfunction than males in many rodent models of diet-induced MetS [43,44]. Surprisingly, we did not detect strong sex-specific response to mWD feeding, since both males and females had altered glucose metabolism and developed hepatic steatosis to a similar extent. It is well established that estrogens play a critical role in protecting females against metabolic syndrome [45,46]. In our study, juvenile female mice experienced nutritional challenge before the onset of sexual maturation, possibly overriding the protective effects of sex steroids. Therefore, our model provides an interesting opportunity to study metabolic dysfunction in both sexes.

Gut microbiota has emerged as a contributor of metabolic dysfunction development, including insulin resistance, dyslipidemia and hepatic steatosis [4,47]. Interestingly, among children with obesity, those suffering from metabolic dysfunction have specific alterations of the gut microbiota composition [3,48,49]. While previous studies have demonstrated that probiotic treatment can improve the metabolic alterations of obese children [50,51], we did not found beneficial effects of the strain *Lactiplantibacillus plantarum* WJL (LpWJL) in mWD-fed juvenile mice. Positive outcome of treatment with LpWJL have previously been demonstrated upon low-protein low-fat feeding in juvenile mice [20], emphasizing the strong connection between probiotic effectiveness and nutritional context. Future investigations will be required to identify bacterial strains or consortia which could target metabolic dysfunctions induced by specific alterations in nutritional environment upon mWD, and the mechanisms underlying these effects.

In conclusion, our study shows that feeding juvenile mice a diet that recapitulates western dietary intake combined with moderate protein restriction induces metabolic dysregulation without obesity, independently of sex. The results of this study might be used to model metabolic alterations in young children and gain new insights in the molecular mechanisms underpinning the development of this understudied pathology.

## Acknowledgements

The authors thank the animal facility of IGFL (PEHR) where animal experiments were conducted, and the associated group for animal welfare (SBEA). They also acknowledge Héloïse Dusson-Andreu for her help in setting up liver histology. The authors would like to thank Antonio Molinaro (University of Gothenburg) for his insightful comments of the manuscript.

## Author contributions

AJ, FDV and FL conceived and designed research. AJ, JLT, AL and EC performed experiments. AJ and EC analyzed data. AJ and FDV interpreted results of experiments. AJ drafted manuscript. AJ, FDV and FL edited and revised manuscript. All authors approved final version of manuscript.

## Availability of data and materials

The dataset supporting the conclusions of this article are included within the article and its additional files.

---

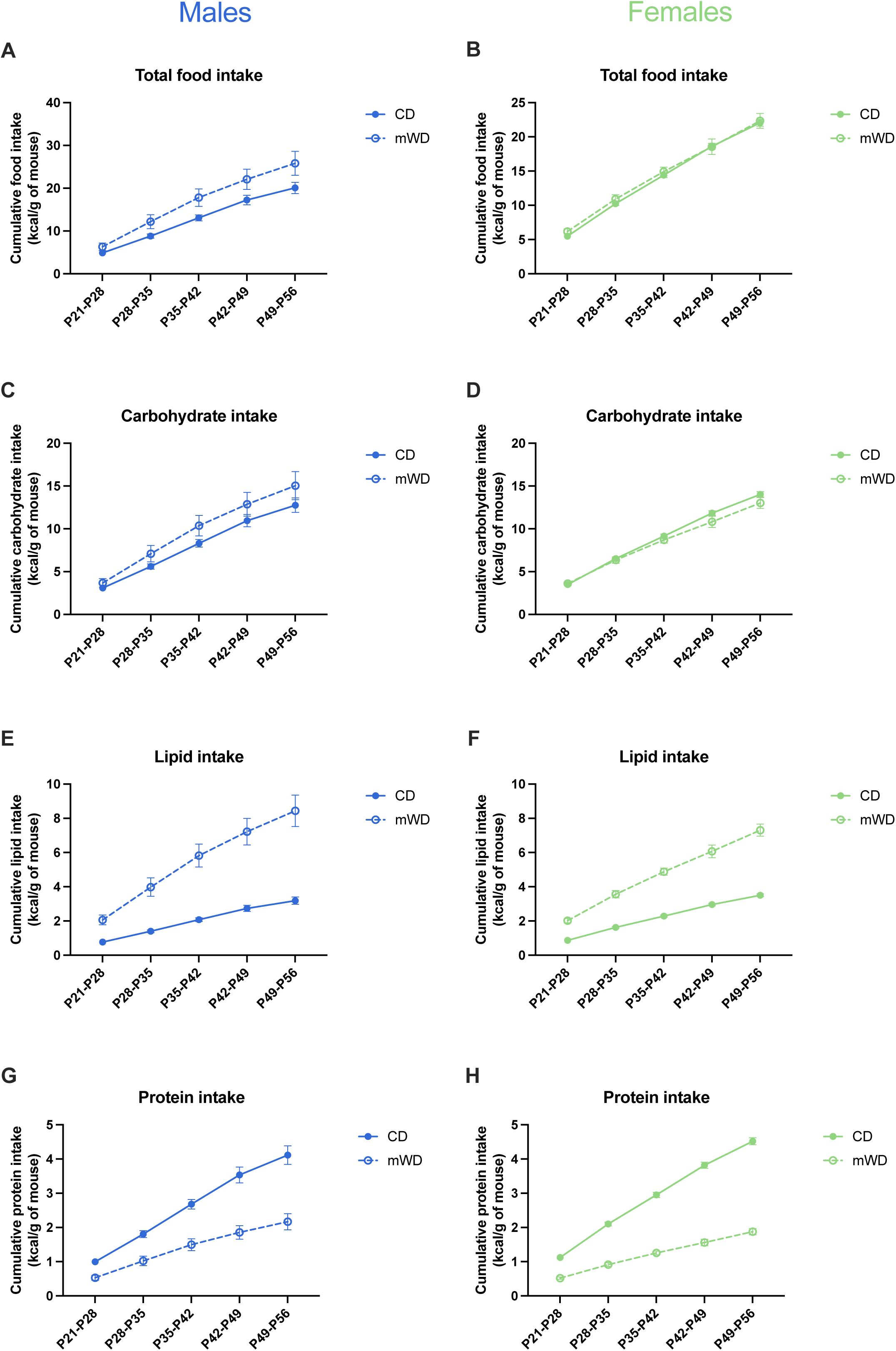

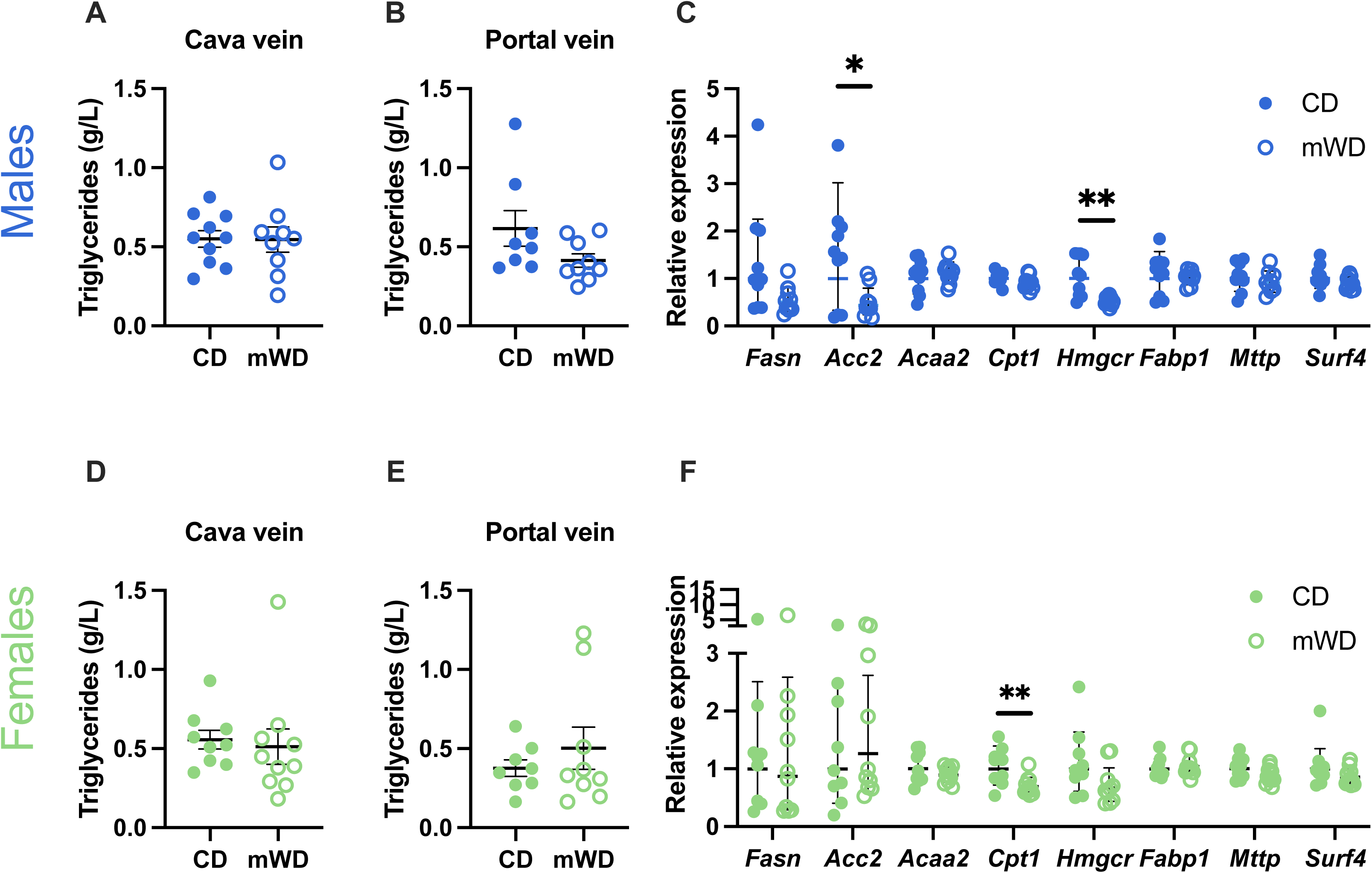

