## Supplementary material for "Protein restriction associated with high fat induces metabolic dysregulation without obesity in juvenile mice"

Supplementary figure legends

Figure S1: mWD-fed animals consume more lipids and less protein than CD-fed animals.

Cumulative food (A-B), carbohydrate (C-D), lipid (E-F) and protein (G-H) intake of male (blue) and female (green) animals fed CD or mWD.

Data are expressed as mean ± SEM.

Figure S2: Circulating triglycerides and hepatic expression of lipid metabolism enzymes are not affected by mWD feeding.

Triglyceride concentration in the serum from cava vein (A, D) and portal vein (B, E) of male (blue) and female (green) animals fed CD or mWD. (C, F): Liver *Fasn*, *Acc2*, *Acaa2*, *Cpt1*, *Hmgcr*, *Fabp1*, *Mttp* and *Surf4* expression in CD and mWD-fed males and females.

(A, B, D, E): Data are expressed as mean ± SEM. (C, F): Data are expressed as geometric mean ± geometric SD.

* p<0.05, ** p<0.01, unpaired t-test.

|  |  | Control Diet (AIN93G) | Modified western diet (mWD) |
| --- | --- | --- | --- |
| **Carbohydrates (g/kg)** | Corn starch | 397.486 | 284.022 |
|  | Maltodextrin | 132 | 80 |
|  | Sucrose | 100 | 286 |
|  | Cellulose | 50 | 50 |
|  | *kcal/g* | *2.52* | *2.60* |
|  | *kcal (% of total)* | *63.6* | *58.3* |
| **Protein (g/kg)** | Casein | 200 | 92 |
|  | L-cystine | 3 | 1.4 |
|  | *kcal/g* | *0.81* | *0.37* |
|  | *kcal (% of total)* | *20.5* | *8.3* |
| **Fat (g/kg)** | Soybean oil | 70 | 31.4 |
|  | Anhydrous milk fat | 0 | 61.1 |
|  | Olive oil | 0 | 28 |
|  | Lard | 0 | 28 |
|  | Corn oil | 0 | 16.5 |
|  | Cholesterol | 0 | 0.4 |
|  | *kcal/g* | *0.63* | *1.49* |
|  | *kcal (% of total)* | *15.9* | *33.4* |
| **Energy density (kcal/g)** |  | **3.96** | **4.46** |
|  | Mineral mix (AIN93G-MX) (g/kg) | 35 | 13.4* |
|  | Vitamin mix (AIN93G-VX) (g/kg) | 10 | 10 |
|  | Choline bitartrate (g/kg) | 2.5 | 2.5 |
|  | TBHQ, antioxidant (g/kg) | 0.014 | 0.028 |
|  | Calcium phosphate, dibasic |  | 10.5 |
|  | Calcium carbonate |  | 6.875 |

*Without Ca &P

Table S1: Composition of the different diets used in the study

| *Gene* | Forward | Reverse |
| --- | --- | --- |
| *Col1a2* | AGTCGATGGCTGCTCCAAAA | AGCACCACCAATGTCCAGAG |
| *Col3a1* | TGGCACAGCAGTCCAACGTA | CATAGGACTGACCAAGGTGGCT |
| *Col4a1* | GGCCCTTCATTAGCAGGTGT | GTGAGGACCAACCGTTAGGG |
| *Des* | GGCCTTGGATGTGGAGATCG | TAGCCTCGCTGACAACCTCT |
| *Vim* | AGACCAGAGATGGACAGGTGA | TTGCGCTCCTGAAAAACTGC |
| *Fasn* | GGAGGTGGTGATAGCCGGTAT | TGGGTAATCCATAGAGCCCAG |
| *Acaa2* | CCTGCTACGAGGTGTGTTCA | GAAGTCCTTGAGAAGGCCCC |
| *Acc2* | CCTTTGGCAACAAGCAAGGTA | AGTCGTACACATAGGTGGTCC |
| *Fabp1* | ATGAACTTCTCCGGCAAGTACC | CTGACACCCCCTTGATGTCC |
| *Mttp* | CTCTTGGCAGTGCTTTTTCTCT | GAGCTTGTATAGCCGCTCATT |
| *Hmgcr* | AGCTTGCCCGAATTGTATGTG | TCTGTTGTGAACCATGTGACTTC |
| *Surf4* | ATGGGACAGAACGACCTGATG | GGTGTCGATATAGTCACGCTG |
| *Cpt1* | CTCCGCCTGAGCCATGAAG | CACCAGTGATGATGCCATTCT |
| *Rpl32* | CCTCTGGTGAAGCCCAAGATC | TCTGGGTTTCCGCCAGTTT |
| *Actb* | GGCTGTATTCCCCTCCATCG | CCAGTTGGTAACAATGCCATGT |
| *Tbp* | CAAACCCAGAATTGTTCTCCTT | ATGTGGTCTTCCTGAATC |

Table S2: Primers used for qPCR
